# Accessibility to sequential working memory fluctuates unconsciously in a theta phase-dependent manner

**DOI:** 10.1101/2021.03.18.435754

**Authors:** Takuya Ideriha, Junichi Ushiyama

## Abstract

Working memory is active short-term memory storage that is easily accessible for later utilization. There is emerging evidence that memorized items are represented rhythmically on the specific phase of the theta-band (4–7 Hz) neural oscillation. However, it is still unknown how this process impacts the accessibility to the active memory storage. Here we show that simply memorizing sequential information causes theta-band fluctuation in our behaviour (i.e., reaction time, RT). We measured RTs to a visual probe that appeared at sequentially memorized locations after a random interval. Consequently, RTs to the probes fluctuated in the theta range as a function of the random interval, and the behavioural rhythmicity supported the hypothesis of the phase-dependent coding of sequential working memory. The current results demonstrate that our behaviour fluctuates unconsciously in the theta-range when recalling sequential memory, suggesting that accessibility to sequential working memory is rhythmic rather than stable, possibly reflecting theta-phase dependent coding.

---

Working memory is active short-term memory storage that is easily accessible and underlies various activities, such as maintaining phone numbers in mind for a short period^1,2^. This cognitive function is known to be related to various types of goal-oriented behaviour^3^, academic success^4^, and a variety of developmental disorders such as reading disability^5^, attention-deficit/hyperactivity disorder^6^ and autism spectrum disorders^7^.

The Lisman/Idiart/Jensen (LIJ) model is an influential hypothesis explaining the mechanisms of sequential working memory^8,9^ (see Fig. 2f for illustration). According to this model, a piece of information is represented by a neural circuit firing in a gamma (>30 Hz) cycle. When maintaining multiple items, this firing is aligned by theta-gamma “phase coding”^8,10–12^. That is, each gamma cycle is assumed to be locked to a specific phase of theta oscillations and the sequence of gamma cycles represents the sequential order. Previous studies have provided experimental evidence supporting this model in monkeys^13^ and humans^10,14–16^. For example, Wolinski et al.^15^ demonstrated that working memory capacity (i.e., the number of items that can be maintained in working memory) was increased by slower theta-band (4 Hz) electrical stimulation around the parietal lobe, but decreased by faster theta-band (7 Hz) stimulation. Because the LIJ model implies that slower theta activity carries more gamma cycles, this experimental finding is consistent with the model. In addition, previous studies of other cognitive functions, such as spatial navigation^17–19^ and episodic memory^20^, have also discussed whether and/or how phase coding is essential for processing sequential information.

However, little is known about how rhythmic memory representations affect our behaviour when we recall these sequential items from our nervous system. In the current study, using novel psychophysical experiments inspired by a series of previous studies in the field of attention^21–23^, we demonstrated that simply maintaining sequential information led to theta-band rhythmic fluctuation in a behavioural index, reaction time (RT). Importantly, the rhythms of unconscious RT fluctuation corresponded to the rhythm of phase coding, providing behavioural support for the notion that sequential working memory is realized by phase coding in the nervous system.

## RESULTS

### Main experiment. Theta Phase-Dependent Rhythmicity in RT During Sequential Working Memory

To examine the effects of sequential working memory on temporal variation of behaviour, we examined fluctuation of RT to sequentially memorized locations as a function of the length of random intervals between the stimuli and the probe (Fig. 1). Nineteen healthy young volunteers participated in Experiment 1. Data from two participants were removed due to excessively low performance (see Methods). Thus, data from 17 participants were included in the following analyses. As shown in Fig. 1, two red dots appeared sequentially on the screen for 200 ms each, and participants memorized the locations and their sequential order. Importantly, the interval after presenting these two stimuli was randomized between 500 and 1500 ms, making it possible to infer the temporal variation of RT. This procedure was developed on the basis of a series of studies on the neural mechanisms of attention^22–24^. After the random interval, a green dot appeared at either of the memorized locations. If the green dot was in the same location as the first red dot, participants were instructed to press the ] key on a JIS keyboard layout (corresponding to the location of the “ key on a US keyboard layout), whereas, if it was in the same location as the second red dot, participants were instructed to press the [ key on a JIS keyboard (in the same position as the [ key on a US keyboard) with their right index finger. Each set comprised 80 trials, and participants carried out five sets, resulting in 400 trials. Although there was no significant difference in the hit rate (mean ± SD) between the first and second locations (first, 97.6 ± 2.2%; second, 97.4 ± 2.4%; t_17_ = 0.93, p = 0.3682) (Fig. 1b), the RT to the second location was significantly longer than for the first location (first, 564.7 ± 171.9 ms; second, 633.0 ± 244.5 ms; t_17_ = −2.57, p = 0.0206) (Fig. 1c).

**Fig. 1.**
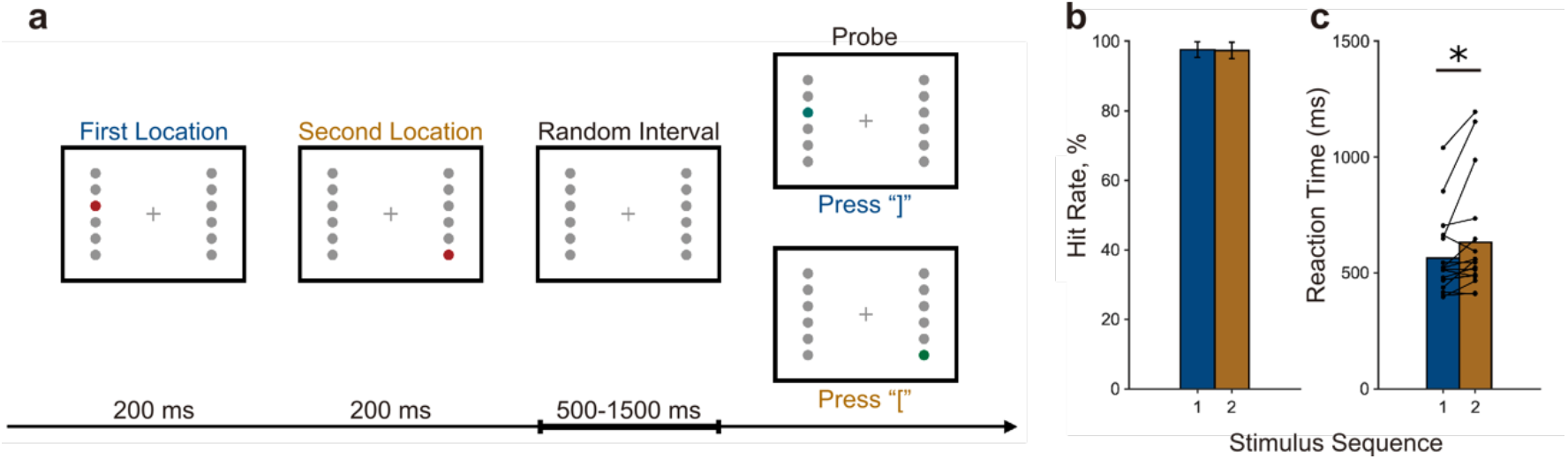
Task Procedure of Experiment 1. (a) Task Procedure. In every trial, two red dots appeared on the screen sequentially and participants memorized the locations and their sequence. After a random interval between 500 and 1500 ms, a green probe dot appeared on either side of the memorized locations. If the probe was in the location of the first red dot, participants were required to press the ] key on a JIS keyboard (corresponding to the “ key on a US keyboard). If it was in the second location, they pressed the [ key on a JIS keyboard (the same position as on a US keyboard) as quickly as possible. The reaction time (RT) to the probe was recorded. (b) Group data (mean ± SD) for hit rate (percentage of correct answers). There was no significant difference in hit rate between the first and second locations (first, 97.6 ± 2.2%; second, 97.4 ± 2.4%; t_17_ = 0.93, p = 0.3682). (c) Group and individual data for reaction time (RT). RTs to the second location were significantly longer than those to the first location (first, 564.6 ± 171.9 ms; second, 633.0 ± 244.5 ms; t_17_ = −2.57, p = 0.0206).

The example distributions of RT as a function of the random interval are shown in Fig. 2a. When we plotted the time course of RT normalized by removing outliers, subtracting the long-term fluctuation of RT due to the effect of learning and variation of concentration^25^, and yielding z-scores (see Methods and Supplementary Fig.1), we found rhythmic fluctuations in RTs as a function of the random interval. In a typical individual, the time courses for each location fluctuated in the theta range as a function of the random interval (Fig. 2b). When the phase relationship between the time courses of RT was mapped, we observed a cluster of data-points in theta range (4–7 Hz) at approximately 270° (Fig. 2c). The 270° phase difference corresponded to the rhythm of phase coding, as illustrated in Fig. 2f. After we performed this analysis for every participant and averaged them, we again observed a clear cluster in the theta range at approximately 270° (Fig. 2d). When we took the peak phase in this analysis for each participant in the theta range and plotted them on a circular histogram, there was a significant tendency toward approximately 270° (260.6° ± 52.2°; Rayleigh test for non-uniformity, p = 0.0071) (Fig. 2e).

**Fig. 2.**
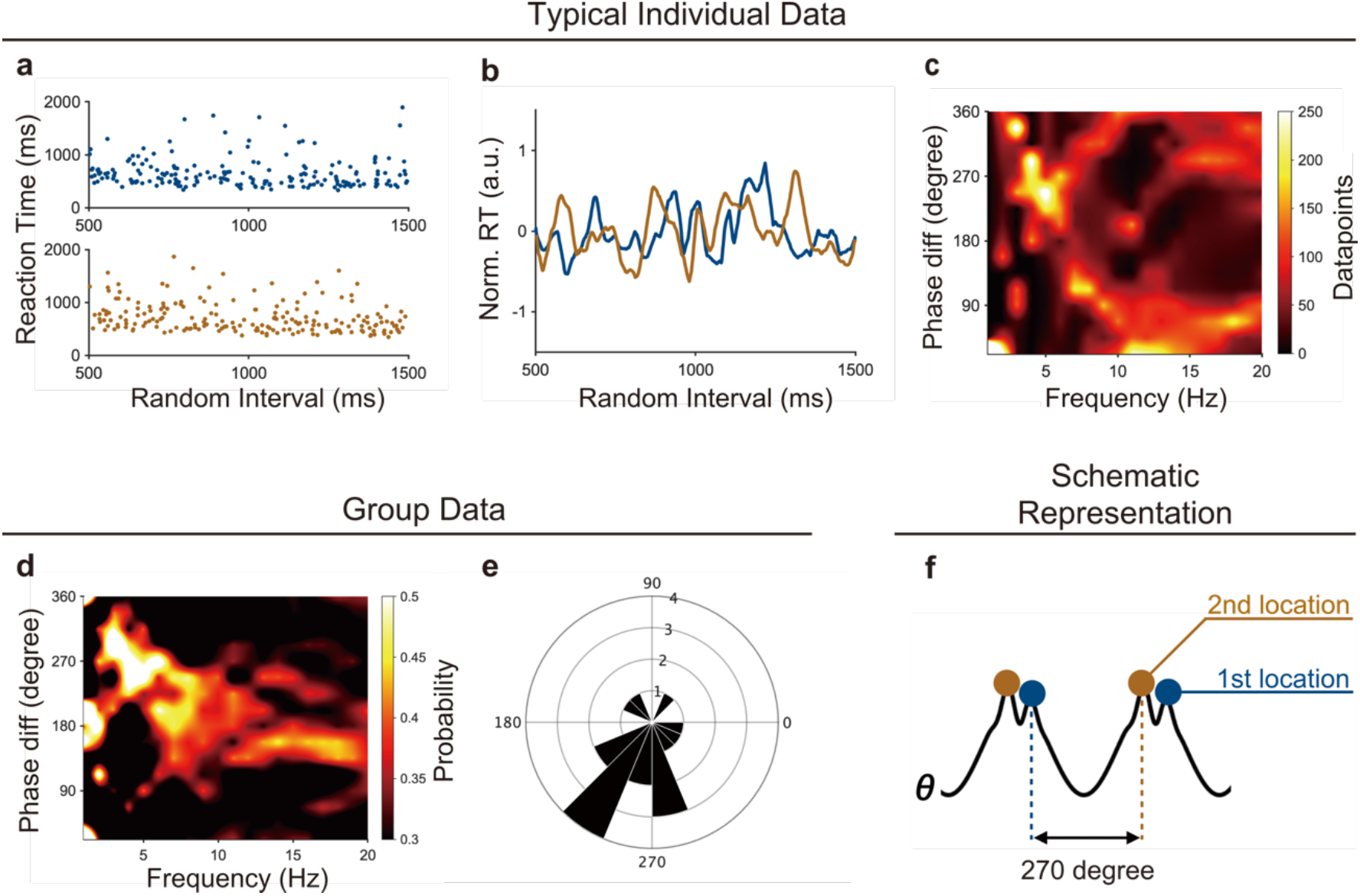
Typical and Group Data for Experiment 1. (a) Example distributions of RTs from a participant. Blue and brown dots represent the RTs to the first and second locations, respectively. Each dot represents RT in one trial. RTs were not uniform, but varied as a function of the randomly presented interval. These distributions were normalized and smoothed to extract RT time courses for each participant (see Methods), then used in the subsequent analyses. (b) Typical data for normalized time courses of RT (Norm. RT). We used data from the same participant shown in (a). The blue waveform denotes the normalized RT time course to the first memorized locations, whereas the brown waveform denotes that of the second memorized locations. We clearly observed the phase shift between the time courses of RTs to the first and second memorized locations. (c) Typical phase difference distribution between time courses of RT to the first and second locations for each frequency. We used data from the same participant shown in (a). Warmer colour indicates a greater number of data-points corresponding to the phase difference. We observed a cluster at approximately 270° around 4–7 Hz (theta-range). (d) Grand average of the phase difference distribution between the two time courses of RT across participants. The mapping was calculated for each participant in the same way as (b), then averaged. Again, we observed a cluster indicating the existence of a phase shift between the two time courses of RT. (e) Circular histogram for the peak of phase differences in the theta (4–7 Hz) range for all participants. The radius denotes the number of participants. The peak phase difference clustered around 270° (256.1° ± 55.7°). This distribution was significantly different from the uniform distribution (Rayleigh test; p = 0.0071). (f) A schematic illustration of the neural oscillation representing two memorized locations suggested by the 270° phase difference. The blue dots represent neural firing for the first memorized location and the brown dots represent neural firing for the second memorized location. Neural firing representing each location appeared to be locked to the specific phase of theta oscillation and their sequence.

To further confirm these results, we conducted another analysis in which we grand-averaged the above-presented time courses of the normalized RTs across participants, based on previous studies^21,22^. This analysis depended on the implicit premise that the rhythmicity of RT fluctuation was at a similar frequency across participants, because the rhythmicity would be canceled by grand-averaging if its frequency varied among individuals. Therefore, if obtained results were similar to the data shown in Fig. 2b-d, it would imply commonality of the oscillatory characteristics such as frequency and phase across participants. Similarly to Fig. 2, we observed rhythmic fluctuations around theta range of RT as a function of the random interval in the grand-averaged data, and the phases were shifted in relation to each other (Fig. 3a). When the phase difference was mapped, a clear tendency toward between 180° and 270° in the theta range was observed (198.6° ± 62.3°; Rayleigh test, p < 0.001) (Fig. 3b). Additionally, a similar but slightly different analysis based on a previous study^23^ was conducted in which we did not subtract the long-term fluctuation of RT but used data only from the third to fifth sets to avoid the effects of learning. Consistently, this additional analysis provided similar results (see Methods and Supplementary Fig.2). Such RT fluctuation was also observed when the duration of visual stimuli was set at 300 ms rather than 200 ms (n = 10, Supplementary Fig.3). Therefore, it was confirmed that the phase relationship of RT fluctuation was irrelevant to the duration of visual stimuli. By adding such grand-averaging analyses, we obtained stronger confirmation that sequential working memory generates rhythmic fluctuation of RT with a similar frequency across individuals.

**Fig. 3.**
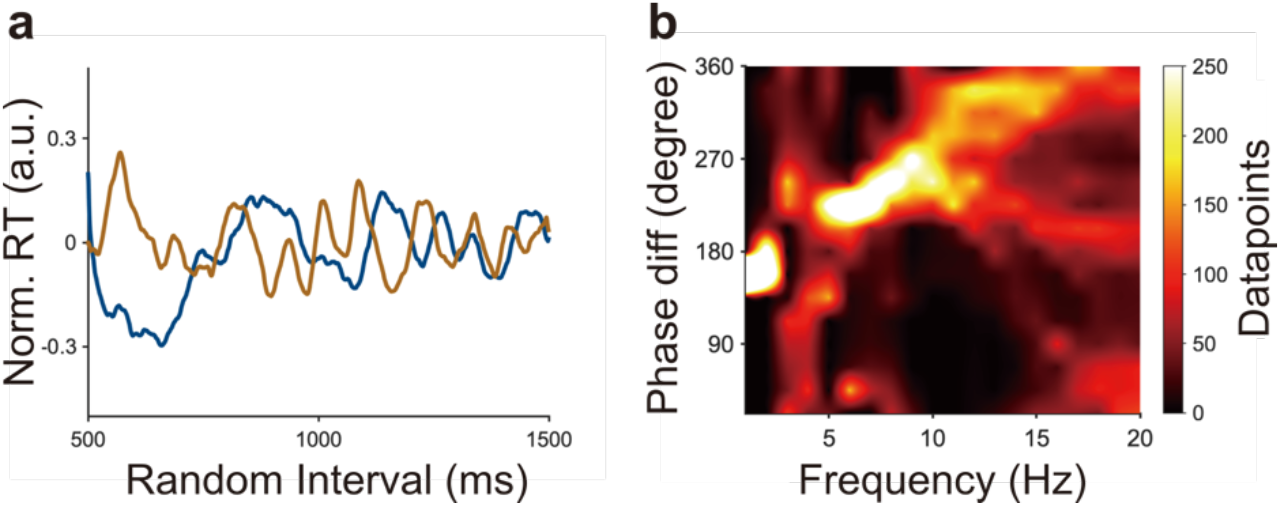
Grand-Averaged Data for Experiment 1. (a) Grand-averaged time courses of normalized RT to the first locations (blue line) and second locations (brown line) across all participants. Fluctuation around the theta range was visible and the two time courses were out of phase, suggesting that accessibility to sequentially memorized items fluctuated in a theta phase-dependent manner. (b) The phase relationship between the two grand-averaged time courses of (a) for each frequency. A clear peak between 180° and 270° in the theta range is visible. Note that this mapping is different from that shown in Fig. 2c, because we obtained this phase relationship after grand-averaging the time courses of RT across all participants.

### Control Experiments

Although the results of Experiment 1 supported the effects of sequential working memory on RT, it remained unclear whether these results were caused by mere reverberation of neural oscillation responding to sensory stimuli^26^, rather than fluctuation due to top-down signals for maintaining working memory items. In addition, it remained unclear whether fluctuation of RT was generated by spatial attention or spatial memory alone, without memorizing sequential information. To examine these possibilities, we conducted the following two control experiments.

First, to examine whether the behavioural rhythmicity obtained in Experiment 1 was due to mere reverberation of neural oscillation to stimuli or spatial attention, we conducted Experiment 2. Fourteen healthy young volunteers participated in this experiment. Unlike Experiment 1, the red dots did not disappear after presentation, but decreased in brightness and remained on the screen. After a random interval between 500 and 1500 ms, a green dot appeared at either location. Participants were instructed to pay attention to the two red dots, and to press the ] key on presentation of the green dot as quickly as possible, regardless of their sequential order.

Using the same analysis applied in Experiment 1, we did not observe a phase relationship similar to that in Experiment 1 (Fig. 4a, b). Note that the 270° peak in the 6–7 Hz range in (Fig. 4a) was due to data from only a few participants. When we plotted peak phase differences for all participants in the theta band on a circular histogram, the distribution was not significantly different from the uniform distribution (Rayleigh test, p = 0.4069) (Fig. 4b). In addition, we obtained similar results in the same analysis as that shown in Fig. 3, in which we grand-averaged the time courses across participants, then mapped the phase relationship (Supplementary Fig.4a, b). These results indicate that the RT fluctuation found in Experiment 1 was not observed in a task requiring only spatial attention.

**Fig. 4.**
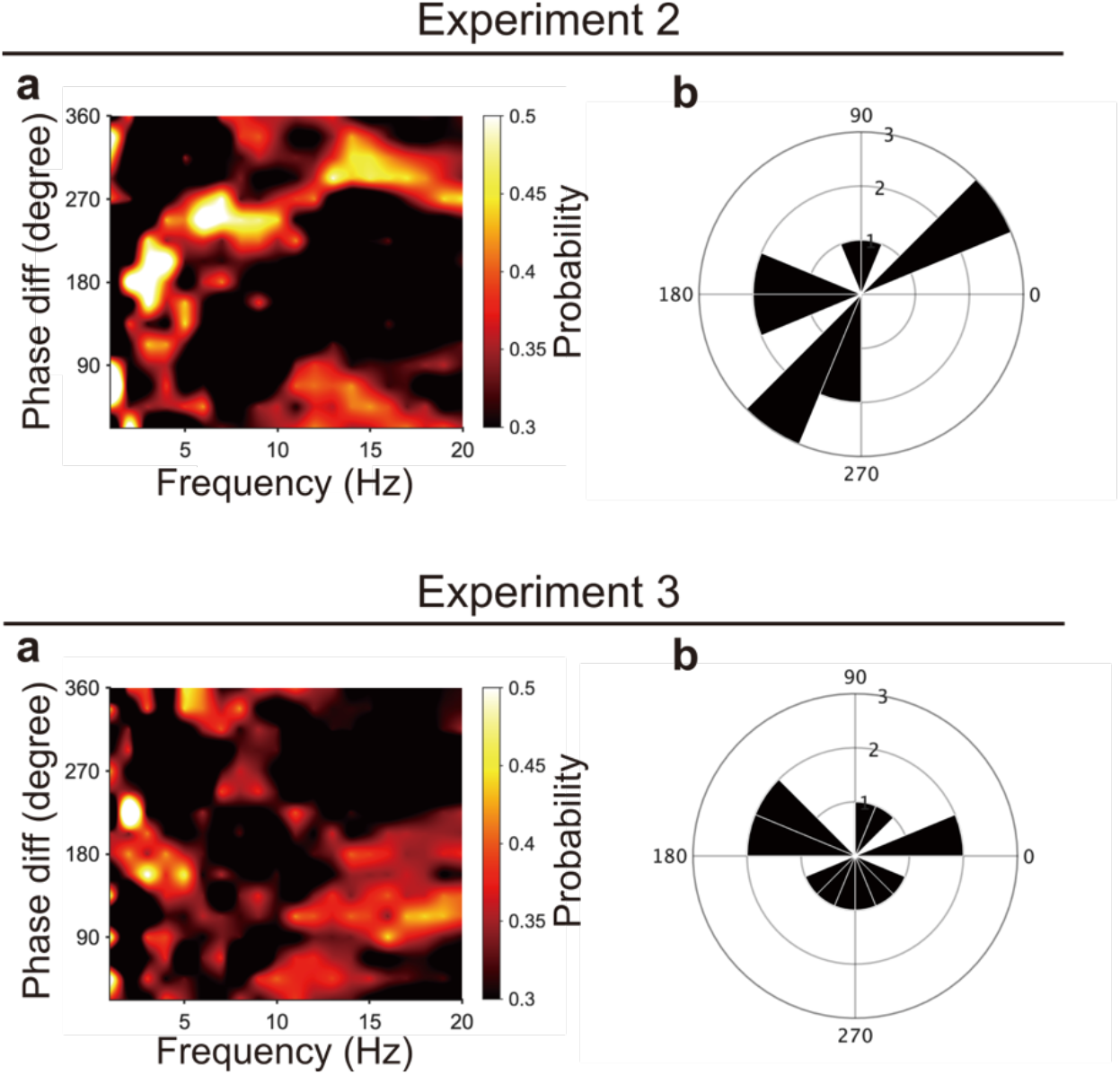
Group Data for Experiments 2 and 3. In Experiment 2, the visual presentation in the task was the same as that in Experiment 1, except that the two red dots remained on the screen, and the task instruction was to attend to the two dots and press the ] key when the colour of either dot changed to green. In Experiment 3, the visual presentation was identical to that in Experiment 1, and the task instruction was to press the ] key when the colour of either dot changed to green. (a and c) Grand-averaged phase relationship between time courses of RT to the first and second stimuli for each frequency across participants for Experiment 2 (a) and Experiment 3 (c). No clear cluster similar to that of Experiment 1 was observed. Note that the 270° peak in the 6–7 Hz range in (a) was due to data from only a few participants. (b and d) The peak position of the phase difference between the two time courses of RT in the theta (4–7 Hz) range for all participants in Experiment 2 (b) and Experiment 3 (d). The radius denotes the number of participants. These distributions did not differ significantly from the uniform distribution (Rayleigh test; B, p = 0.4069; D, p = 0.8771).

Second, we conducted Experiment 3 to examine whether the behavioural rhythmicity obtained in Experiment 1 was caused by spatial working memory without sequential information. Fourteen healthy young volunteers participated in this experiment. In this experiment, all procedures were the same as those in Experiment 2, except that the red dots disappeared from the screen after presentation. That is, the presentation on the screen was identical to that in Experiment 1. Participants were required to memorize the locations of the red dots without memorizing the sequence. In this experiment, participants were instructed to memorize the locations, attend to them, and press the ] key when the green probe dot appeared on the screen. Notably, the average RT in Experiment 3 was marginally faster than that in Experiment 2 (373.0 ± 48.1 ms in Experiment 2; 345.3 ± 39.3 ms in Experiment 3; t_12_ = 1.64, p = 0.1126). Thus, it would be reasonable to assume that participants actually made an effort to memorize the two locations in Experiment 3, compared with Experiment 2 where they just paid attention to the stimuli.

Similarly to the results of Experiment 2, we did not observe a 270° phase relationship (Fig. 4c and Supplementary Fig. 4c, d). The distribution of peak phase differences for all participants was not significantly different from a uniform distribution (Rayleigh test, p = 0.8771) (Fig. 4d). Therefore, fluctuation of RT was not found in a task requiring spatial working memory without maintaining sequential information.

## DISCUSSION

The present study demonstrated that RT fluctuates unconsciously in the theta range when recalling sequential items. This finding suggests that accessibility to sequential working memory is rhythmic rather than stable, in a theta phase-dependent manner. The cause of this RT fluctuation is likely to be the rhythmic nature of neuronal activity representing memorized items. In support of this notion, RT to a stimulus has long been proposed to fluctuate in accordance with the phase of neural oscillation^27,28^. Since highly excited neural populations tend to react quickly to external input^29^, RT fluctuation observed in the current study may be the outcome of the rhythmic excitation of neural circuits representing memorized items.

Interestingly, the observed 270° phase difference in RT fluctuation to the first and second memorized locations corresponded to the rhythm of phase coding, supporting the LIJ model of working memory^8,9^. This model theoretically proposes that, when memorizing sequential items, each gamma cycle representing each item is locked to a specific phase of theta oscillation and the sequence of gamma cycles is considered to represent the order of memorized items. Here, we showed that a 270° phase difference was observed only when participants memorized sequential information, but not when they just paid attention to or maintained spatial locations without a sequential order. These results suggest that items in sequential working memory is represented by phase coding based on the LIJ model, and that its intrinsic fluctuation is strong enough to generate behavioural fluctuation.

Some might doubt that the RT fluctuation would just reflect mere reverberation of cell populations responding to respective stimuli presented for 200 ms each. Such phenomena were reported in the context of neural entrainment to sensory stimuli. Indeed, neural entrainment was observed not only when the stimuli are flickering over a few seconds^30^ but also when only two consecutive stimulation is applied^26^. If the results obtained from Experiment 1 were due to the neural entrainment to rhythmic presentation of visual stimuli, the observed RT rhythmicity would be changed with different duration of visual stimuli. However, when the duration of visual stimuli was set at 300 ms rather than 200 ms, the similar phase-frequency relationship was also obtained (Supplementary Fig.3). Therefore, the rhythmicity of RT fluctuation cannot be explained with the neural entrainment to visual presentation. Additionally, in Experiment 2 in which participants just paid attention to the two locations without memorizing sequential information, the 270° phase relationship was not observed. Moreover, in Experiment 3 in which the visual presentation was completely the same as Experiment 1 but participants were just required to remember spatial locations without sequential order, the phase relationship was not observed, again. Hence, the obtained rhythmicity should not be due to mere reverberation of cell populations.

The results of Experiment 1 revealed an approximately 270° phase difference between the rhythmicity of RTs to the first and second memorized locations. This implies that the neural circuit of the second memorized item was activated 90° earlier than that of the first memorized item on theta oscillation (Fig. 2f). That is, sequentially memorized items appear to be replayed in a backward order. This “backward replay” might seem counterintuitive, but could be a basic mechanism of representing sequential information in the nervous system. In accord with this notion, previous studies reported that after rats explored a space, hippocampal neurons encoding places were found to fire in a temporally reversed order^17,18^, possibly reflecting recollection of past experiences. In addition, in humans, evidence of backward replay has been accumulating in the context of spatial navigation^19^ and working memory^31^. The current results provided complementary evidence of backward replay in relation to theta phase coding.

From the perspective of attention, the current results deepen the understanding of the relationship between attention and working memory^32–36^. Importantly, the design of the present experiment is similar to that of previous studies on the rhythmic theory of attention, reporting that even sustained attention is not static, but fluctuates in the theta to alpha range^21–23,37–39^. For example, Landau and Fries^22^ measured the detection rate of a subtle flash presented randomly in the right or left hemifield after a random interval below 1000 ms. In this task, participants were required to attend to both the right and left hemifield to detect the flash. Consequently, the detection rate of the right and left hemifield each fluctuated around the theta range in anti-phase coordination, suggesting that divided attention to two locations was not stable, but oriented alternately in the theta range. Taking these previous findings together with the current results, maintaining sequential information in mind could potentially “bias” the anti-phase rhythm of attention toward a rhythm with a 270° phase difference in the theta range.

In conclusion, the present findings demonstrated that human behaviour fluctuates unconsciously in the theta range when sequential items are recalled from working memory. This suggests that accessibility to memorized items is rhythmic rather than stable possibly in accordance with theta phase-dependent representation of sequential working memory. In future, behavioural approaches like the current study could be used to complement physiological and/or imaging findings, and advance understanding of the neural basis of cognitive functions and its impact on our behaviour.

## Methods

### Experimental model and participant details

Twenty-Nine individuals participated in Experiment1. Nineteen of them (mean ± SD, 20.4 ± 1.9 years; 10 females, one left handed) carried out the main experiment with 200 ms duration of visual stimuli, while the other 10 participants (21.7 ± 2.8 years; four females, all right handed) carried out the sub experiment with 300 ms duration (See Experimental procedure for the detail). Fourteen individuals (21.0 ± 2.2 years; four females, all right handed) participated in Experiment 2, and 14 individuals (20.6 ± 2.3 years; eight females, one left handed) participated in Experiment 3. In Experiment 1, data from two participants were removed, resulting in 17 participants’ data. One participant was excluded because of an excessively low hit rate (50.5%) which was similar to the level of chance (50%), whereas the other exhibited excessively slow and unstable RTs (mean ± SD, 2260 ± 2090 ms). All participants had normal or corrected-to normal visual acuity. Participants gave informed consent before the experimental session and received monetary compensation. All experimental protocols and procedures were approved by the SFC Research Ethics Committee, Keio University (Approval Number 293).

### Experimental procedure

The stimuli and experimental program were generated using JavaScript (jsPsych toolbox^40^ and jsPsych-Psychophysics plugin^41^). All experiments were conducted online to prevent the spread of COVID-19. Therefore, stimuli were presented on participants’ computers. The validity of online experimental designs has been repeatedly confirmed, including the precision of RT measurement^42,43^. We confirmed that the refresh rate of each participant’s monitor was at least 50 Hz to enable accurate tracking of behavioural oscillations in the theta (4–7 Hz) range. The monitor was positioned approximately 40 cm from the participant. During all experiments, participants were instructed to gaze at a fixation cross at the centre of the screen, except during the inter-trial interval or rest period. Throughout all experiments, the fixation cross and 12 gray (Colour code #c0c0c0) filled circles remained on the screen (Fig. 1a). Six circles lined up vertically on the left side of the screen and the other six lined up on the right. The width between the left and right circles was set to 21° of visual angle, the height between the top and bottom circles was set to 15°, and the circle size was set to 1° × 1°.

In Experiment 1, the presentation of “Trial n (the number of the trial)” indicated the beginning of each trial. After 1000 ms, two red (Colour code: #c40000) dots (1° × 1°) sequentially appeared on the screen for 200 ms each in the main experiment, while the duration was set at 300 ms in the sub experiment. Participants memorized the sequence and locations of the dots. After a random interval of 500–1500 ms, a green (Colour code: #00c462) probe dot (1° × 1°) appeared at either of the locations of the two red dots. The green dot remained on the screen until participants pressed the ] or [ key on a JIS keyboard. These keys correspond to the “ and [ keys on a US keyboard, respectively. If the green dot was in the location of the red dot that appeared first, the correct response was ], and vice versa. Participants were instructed to press the correct key as quickly as possible. Each set included 80 trials, and participants performed five sets, resulting in 400 trials in total. Before the main experiment, a practice session including 40 trials was conducted.

In Experiment 2, the flow of the presentation of stimuli was the same as the main experiment of Experiment 1 except that the red dots did not disappear from the screen and participants did not memorize the sequence of stimuli. Instead, the red dots decreased in brightness (Colour code: #c47676) after 200 ms and remained until participants responded. The requirement of participants was to press the ] key as quickly as possible after the green probe appeared at either of the locations of the two red dots. To achieve this, participants were instructed to direct their attention to the two red dots. The number of sets and trials was the same as in Experiment 1.

In Experiment 3, the flow of the presentation of stimuli was the same as in the main experiment of Experiment 1. The only difference from Experiment 1 was the requirements of the participants. Participants were not required to memorize the order of stimuli, but were instructed to press the ] key when the green probe appeared, regardless of its location. Therefore, participants were required to memorize and direct attention to the location of the red dots and respond as quickly as possible. The number of sets and trials was the same as in Experiment 1.

### Data Analyses

In each experiment, we analyzed RT fluctuation as a function of the length of the random interval. In Experiment 1, only data from correct responses were analyzed, whereas all data were analyzed in Experiment 2 and 3. The analyses described below were conducted individually and later applied to other analyses. Behavioural time courses were analyzed separately for RT to the location of the first and second presented red dot. Hereafter, we refer to the former time course as “first waveform”, and the latter time course as “second waveform”. In the first waveform, trials with RTs longer than 3000 ms or shorter than 200 ms were eliminated from datasets for the following analyses because of an excessive lack of concentration or mistakes. As shown in Supplementary Fig.1, to remove the effects of learning and fluctuations of concentration^25^, we subtracted the lower envelope of the time series from the original RT time course. Subsequently, trials with RT exceeding two standard deviations were removed to exclude outliers. These processes resulted in omitting 6.9 ± 3.5%, 4.6 ± 1.4%, and 4.6 ± 1.9% of trials in Experiment 1, 2 and 3, respectively. Then, to extract behavioural time courses, we sorted the RTs in order of the length of the interval (Fig. 2a), then shifted a 50-ms window in steps of 1 ms from 500 to 1500 ms and re-calculated the averaged RT in each window. Both time courses were smoothed and then z-transformed, yielding time courses at a sampling rate of 1000 Hz. When analyzing averaged time courses across participants, the z-transformed time courses were used for averaging. To examine the reliability of the results from Experiment 1, we replicated the results using other analyses, as follows (Supplementary Fig.2). In an additional analysis^23^, we did not subtract the lower envelope to remove the effects of learning, but used data from the third to fifth sets only with a time window of 100 ms.

To map the phase relationships between the first and the second waveforms for each frequency, we first convolved complex Morlet wavelets (three cycles) with those two waveforms from 1 to 20 Hz (in 1-Hz steps) to extract each frequency component. Then, the time course of the phase difference was calculated by subtracting the angle of the second waveform from that of the first waveform. The 1000 sample points of this time course of the phase difference were divided into 16 bins of angle differences equally distributed from 0 to 360°. The phase difference distributions were mapped for all frequencies from 1 to 20 Hz (e.g., Fig. 2c). Then, the distribution of the peak phase differences between 4 and 7 Hz for all participants were visualized with a circular histogram (e.g., Fig. 2e). The nonuniformity of the phase relation was tested using the Rayleigh test (CircStats toolbox^44^).

## Code availability

The computer code that support the findings of this study are available from the corresponding author upon reasonable request.

## Data availability

The data that support the findings of this study are available from the corresponding author upon request.

## ACKNOWLEDGEMENTS

This work was supported by a designated donation from Living Platform, Ltd, Japan to J.U., Taikichiro Mori Memorial Research Grants to T.I., and Sasakawa Scientific Research Grant (The Japan Science Society, JSS) to T.I. We thank Ms. Tomomi Hamaoka, Ms. Kana Iijima, and Ms. Chieko Matsuda for their secretarial assistance and Mr. Ryoichiro Yamazaki, Mr. Hirotaka Sugino, Ms. Rina Suzuki and all other members of our laboratory for their insightful comments on the work. We thank Benjamin Knight, MSc., from Edanz Group (https://en-author-services.edanz.com/ac) for editing a draft of this manuscript.

## AUTHOR CONTRIBUTIONS

Conceptualization, T.I. and J.U.; Methodology, T.I.; Software Development, T.I.; Data acquisition, T.I.; Data Analyses, T.I.; Visualization, T.I.; Interpretation, T.I. and J.U.; Writing – Original Draft, T.I.; Writing – Review & Editing, T.I. and J.U.; Supervision, J.U.; Funding Acquisition, T.I. and J.U.

