## Supplementary Information for "Accessibility to sequential working memory fluctuates unconsciously in a theta phase-dependent manner"

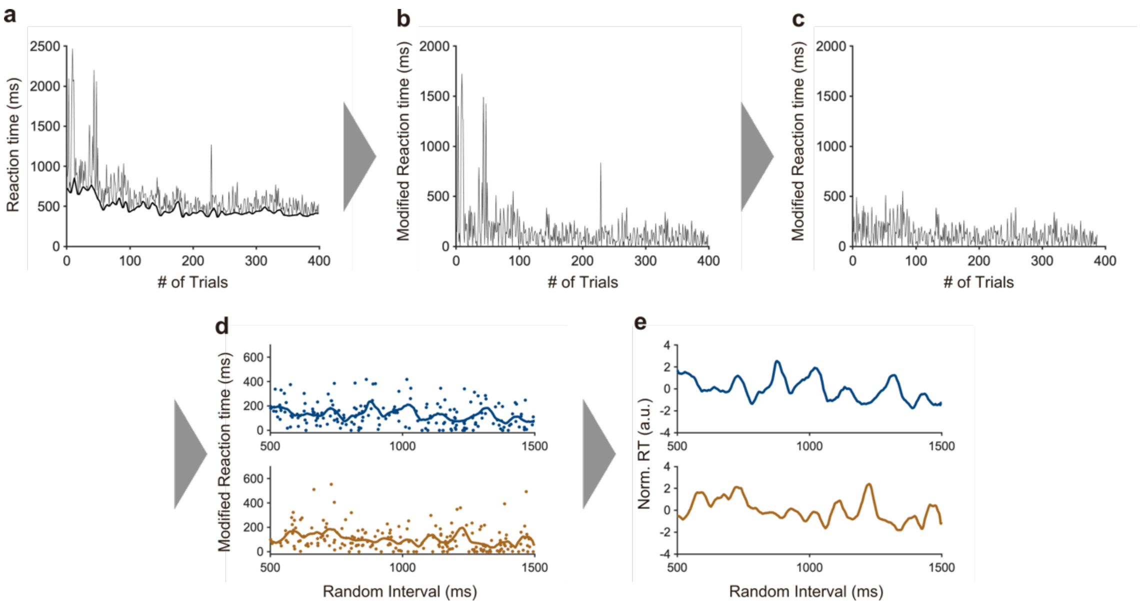


**Supplementary Fig. 1. Preprocessing of RT Time Courses**

(a) RTs fluctuated in the order of tens of seconds due to the effect of learning and variation of concentration. To focus on subtle fluctuations due to neural oscillation, we subtracted lower envelopes from the RT time courses (a to b). After subtracting the envelopes, trials with RTs over two standard deviations from the mean were removed (b to c). The modified RTs were divided into two distributions due to their temporal order and aligned as a function of a random interval to extract the RT waveforms (c to d). Finally, the waveforms were normalized to produce the normalized reaction time (Norm. RT) (d to e).

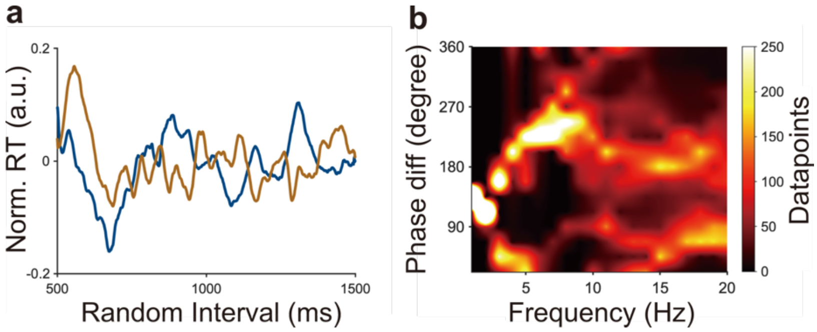


**Supplementary Fig. 2. Additional Analyses in Experiment 1 Related to Fig. 3**

(a) Grand-averaged time courses of RT to the first and second locations across all participants. The blue line denotes the RT time course to the first locations, whereas the brown line denotes the RT time course to the second locations. We did not subtract the lower envelope and used a 100-ms time window to extract the time courses in preprocessing of RTs, but used only the third to fifth sets that appeared to be less affected by the effects of learning.

(b) Phase relationship between the blue and brown time courses of (a) for each frequency. A cluster is visible in the theta range between 180° and 270°.

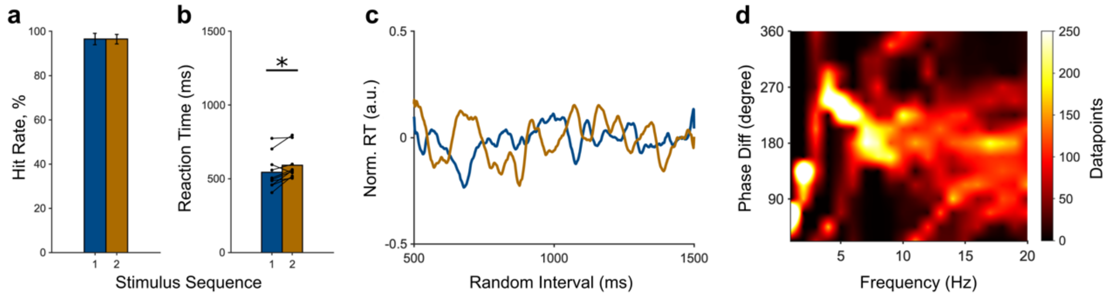


**Supplementary Fig. 3. Results from the sub experiment of Experiment 1**

Similar results were obtained when the duration of visual stimuli was set at 300 ms rather than 300. Ten healthy volunteers completed six sets of 80 trials resulting in 480 trials.

(a) Group data (mean ± SD) for hit rate. There was no significant difference in hit rate between the first and second locations (first, 96.5 ± 2.6%; second, 96.5 ± 2.1%; $t_{10} = 0.15$, $p = 0.8860$).

(b) Group and individual data for RT. RTs to the second location were significantly longer than those to the first location (first, 543.7 ± 116.8 ms; second, 591.5 ± 107.1 ms; $t_{10} = -3.00$, $p = 0.0149$).

(c) Grand-averaged time courses of normalized RT to the first locations (blue line) and second locations (brown line) across all participants. Similar RT fluctuations and the phase relationship are visible to when the duration was 200 ms. This result was obtained through completely the same analysis as Fig. 3a.

(d) The phase relationship between the two grand-averaged time courses of (b) for each frequency. A clear tendency toward between 180° and 270° in the theta range is visible (212.8° ± 54.7°; Rayleigh test, $p < 0.001$). This result was obtained through the same analysis as Fig. 3b.

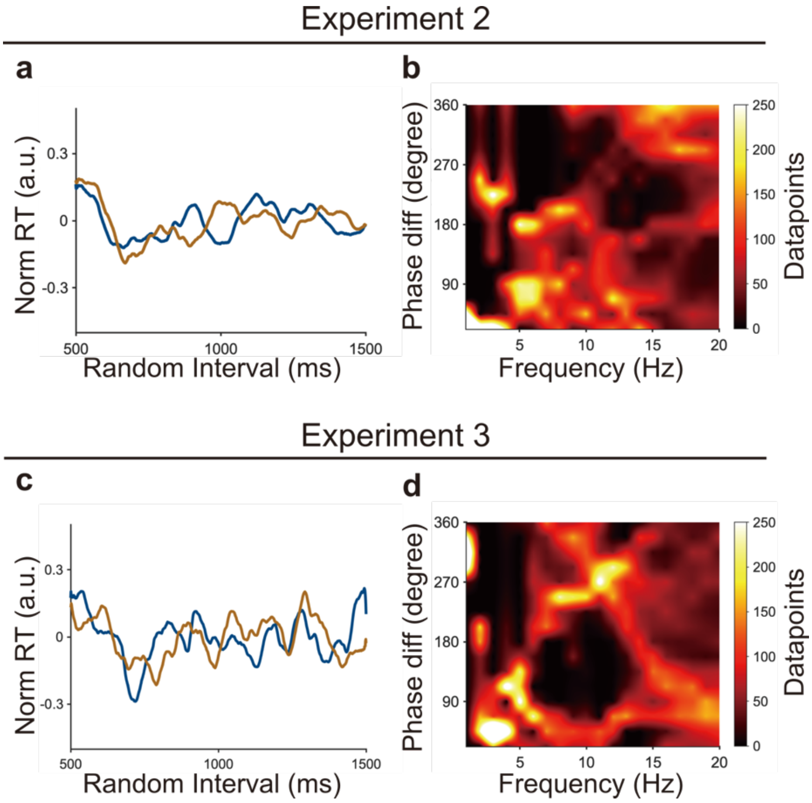


**Supplementary Fig. 4. Grand-Averaged Data for Experiments 2 and 3**

The same analyses shown in Fig. 3 were conducted with the data for Experiments 2 and 3.

(a and c) Grand-averaged time courses of normalized RTs to the first and second locations across all participants for Experiment 2 (a) and Experiment 3 (c). The blue lines are RT time courses to the first locations and the brown lines are RT time courses to the second locations.

(b and d) Phase relationship between the RT time courses to the first and second locations. We did not observe the 270° phase relationship in the theta range.
